# Molecular and pharmacological modulators of the tumor immune contexture revealed by deconvolution of RNA-seq data

**DOI:** 10.1101/223180

**Authors:** Francesca Finotello, Clemens Mayer, Christina Plattner, Gerhard Laschober, Dietmar Rieder, Hubert Hackl, Anne Krogsdam, Zuzana Loncova, Wilfried Posch, Doris Wilflingseder, Sieghart Sopper, Marieke Ijsselsteijn, Douglas Johnson, Yaomin Xu, Yu Wang, Melinda E. Sanders, Monica V. Estrada, Paula Ericsson-Gonzalez, Justin Balko, Noel de Miranda, Zlatko Trajanoski

## Abstract

The immune contexture has a prognostic value in several cancers and the study of its pharmacological modulation could identify drugs acting synergistically with immune checkpoint blockers. However, the quantification of the immune contexture is hampered by the lack of simple and efficient methods. We developed quanTIseq, a deconvolution method that quantifies the densities of ten immune cell types from bulk RNA sequencing data and tissue imaging data. We performed extensive validation using simulated data, flow cytometry data, and immunohistochemistry data from three cancer cohorts.

Analysis of 8,000 samples showed that the activation of the CXCR3/CXCL9 axis, rather than the mutational load is associated with cytotoxic T cell infiltration. We also show the prognostic value of deconvolution-based immunoscore and T cell/B cell score in several solid cancers. Finally, we used quanTIseq to show how kinase inhibitors modulate the immune contexture, and we suggest that it might have predictive value for immunotherapy.

## INTRODUCTION

Cancer immunotherapy with antibodies targeting immune checkpoints has shown durable benefit and even curative potential in various cancers^1,2^. As only a fraction of patients respond to immune checkpoint blockers, efforts are underway to identify predictive markers for cancer immunotherapy and mechanistic rationale for combination therapies. We have previously shown that the immune contexture — the type and density of tumor-infiltrating immune cells — has a prognostic value in colorectal cancer (CRC)^3^. Later, the association between the densities of tumor-infiltrating immune cells and patient overall survival was confirmed in different primary and metastatic cancers^4^. In particular, cytotoxic CD8^+^ T cells, which can specifically recognize and kill tumor cells, have been associated with a good clinical outcome in different cancer types^5^ and are recognized to have a pivotal role in anti-PD1 immunotherapy^1^. Therefore, the quantification of the immune contexture of human tumors cannot only unveil prognostic markers, but can also provide relevant information for the prediction of response to checkpoint blockade.

The quantification of the immune contexture of archived tumor samples holds the promise to identify drugs having additive or synergistic potential with immune checkpoint blockers. For example, since certain chemotherapeutic drugs induce immunogenic cell death^6^, the analysis of large number of samples could pinpoint patient subgroups that would benefit from the combination with immune checkpoint blockers. Similarly, as a number of targeted anticancer agents exhibit immunostimulatory activity^6^, the quantification of the immune contexture could provide mechanistic rationale for the design of combination therapies. However, comprehensive and quantitative immunological characterization of tumors in a large number of clinical samples is currently hampered by the lack of simple and efficient methods. Cutting-edge technologies like single-cell RNA sequencing and multi-parametric flow or mass cytometry are technically and logistically challenging and cannot be applied to archived samples. Multiplexed immunohistochemistry (IHC)^7^ or immunofluorescence (IF) assays can be performed only in specialized labs and require sophisticated equipment and extensive optimization of protocols for specific cancer entities. Moreover, manual and semi-automatic image analysis is required, which is highly time consuming and laborious.

Computational methods for quantitative immunophenotyping of tumors from bulk RNA sequencing (RNA-seq) data hold potential for efficient and low-cost profiling of a large number of samples, but currently suffer from several limitations. Bioinformatics methods based on gene signatures of immune cells, like MCP-counter^8^, xCell^9^, and other approaches based on gene set enrichment analysis (GSEA)^10–12^, compute only scores that predict the enrichment of specific immune cell types^13^ and hence, do not provide quantitative information about cell proportions. Deconvolution algorithms (reviewed in ^14^) enable quantitative estimation of the proportions of the cell types of interest. However, currently available deconvolution algorithms for immune cell quantification have several drawbacks^14^. For instance, CIBERSORT, a popular method based on support-vector regression for the deconvolution of 22 immune-cell phenotypes, can only infer cell fractions relative to the total immune-cell population and has been developed and validated using microarray data^15^. TIMER performs deconvolution of six immune cell types, but the results cannot be interpreted directly as cell fractions, nor compared across different immune cell types and data sets^15^. EPIC, a deconvolution method recently developed using RNA-seq data, estimates absolute fractions referred to the whole cell mixture, but does not consider immune cells relevant for cancer immunology like regulatory T cells (T_reg_) cells, dendritic cells, and classically (M1) and alternatively (M2) activated macrophages^16^. Over and above, these methods have not been independently validated using data from bulk RNA-seq from solid tumors and from IHC- or IF-stainings from the same samples. Hence, there is a need for a validated deconvolution-based algorithm that estimates absolute proportions of relevant immune cell types from RNA-seq data and thereby enabling inter-sample as well as intra-sample comparisons.

We therefore developed quanTIseq, a computational pipeline for the characterization of the tumor immune contexture using bulk RNA-seq data and imaging data from whole tissue slides. quanTIseq can quantify the absolute fractions of immune cells using a novel deconvolution approach and performs *in silico* multiplexed immunodetection of the same cell types by integrating the deconvolution results with total cell densities extracted from images from IF, IHC, or haemotoxylin and eosin (H&E)-stained tissue slides. We performed extensive validation using simulated data, published data sets, and *de novo* generated flow cytometry data. In addition, we validated quanTIseq using RNA-seq data and histological images from IHC/IF stained slides from three independent cancer data sets. We then applied quanTIseq to analyze over 8,000 solid tumors of The Cancer Genome Atlas (TCGA)^17^ and show that the activation of the CXCR3/CXCL9 axis, rather than the mutational load, is associated with the infiltration of intratumoral cytotoxic T cells. Moreover, we observe highly heterogeneous immune contextures across and within tumors, and show that the immunoscore and a T cell/B cell score computed from quanTIseq deconvolution results have prognostic values in several solid cancers. Finally, we show that the immune contexture of human tumors is pharmacologically modulated by kinase inhibitors and show the potential of quanTIseq to identify immune contexture underlying the response to therapy with immune checkpoint blockers.

## RESULTS

### Development of quanTIseq deconvolution algorithm

We developed quanTIseq, a computational pipeline for the analysis of raw RNA-seq and tissue imaging data that quantifies the fractions and densities of ten different immune cell types relevant for cancer immunology (Figure 1a). We first designed a novel signature matrix using RNA-seq data (Figure 1b and **Supplementary Table 2**). To this end, we collected a compendium of 51 publicly available RNA-seq data sets (**Supplementary Table 1**) from ten different immune cell types: B cells, M1 and M2 macrophages, monocytes (Mono), neutrophils (Neu), natural killer (NK) cells, non-regulatory CD4^+^ T cells, CD8^+^ T cells, T_reg_ cells, and dendritic cells (DC). These data were integrated with additional large-scale data resources from immune and non-immune cells and used to select the signature genes with the highest specificity and discriminative power to construct the immune-cell signature matrix (details in **Methods**).

**Figure 1:**
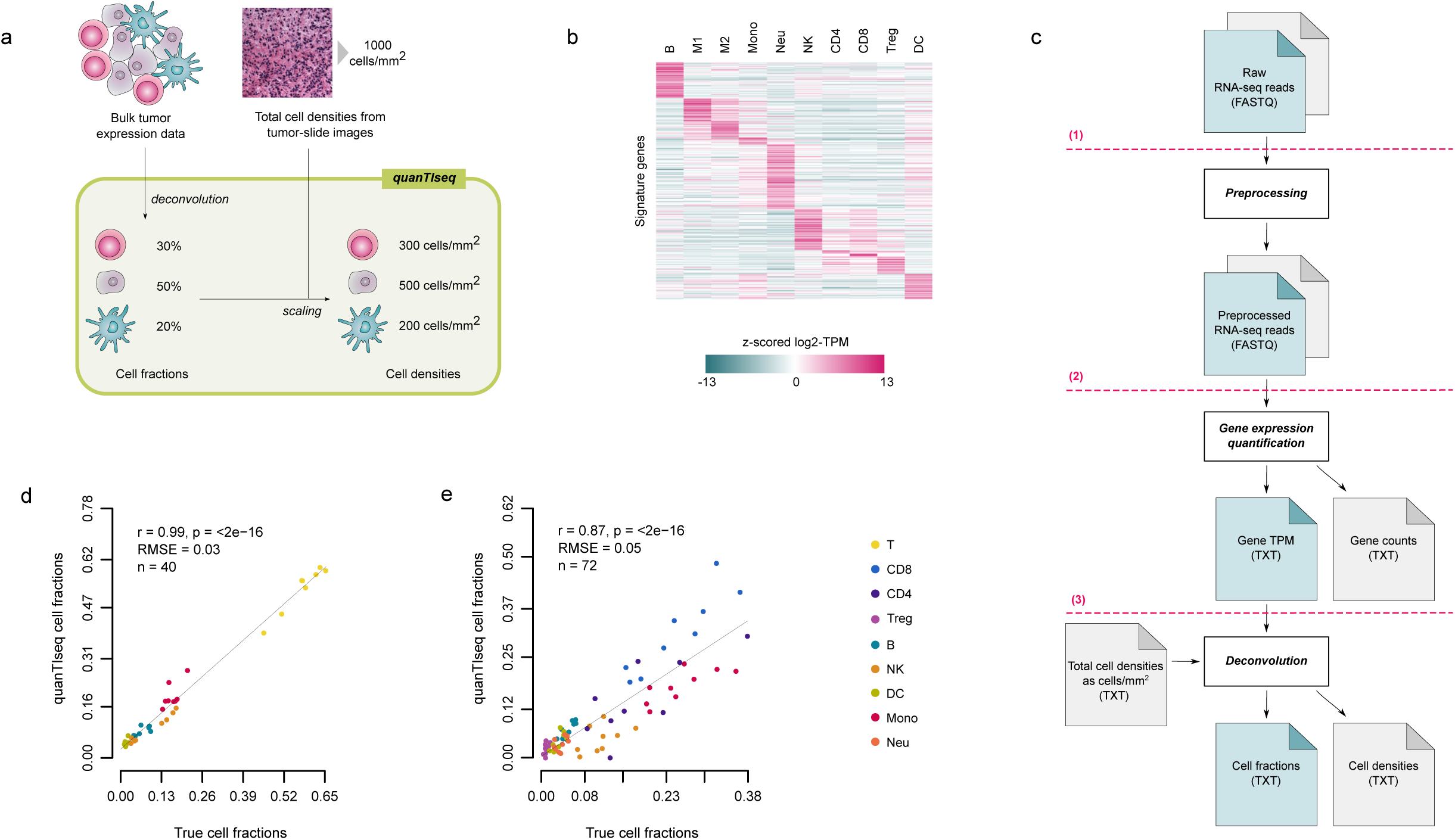
quanTIseq method and validation based on blood-cell mixtures. **(a)** quanTIseq characterizes the immune contexture of human tumors from expression and imaging data. Cell fractions are estimated from expression data and then scaled to cell densities (cells/mm^2^) using total cell densities extracted from imaging data. **(b)** Heatmap of quanTIseq signature matrix, with z-scores computed from log_2_(TPM+1) expression values of the signature genes. **(c)** The quanTIseq pipeline consists of three modules that perform: (1) pre-processing of paired- or single-end RNA-seq reads in FASTQ format; (2) quantification of gene expression in transcripts-per-millions (TPM) and gene counts; (3) deconvolution of cell fractions and scaling to cell densities considering total cells per mm^2^ derived from imaging data. The analysis can be initiated at any step. Optional files are shown in grey. Validation of quanTIseq with RNA-seq data from blood-derived immune-cell mixtures generated in ^23^ **(d)** and in this study **(e)**. Deconvolution performance was assessed with Pearson’s correlation (r) and root-mean-square error (RMSE) using the flow cytometry estimates as ground truth. The line represents the linear fit. B: B cells; CD4: non-regulatory CD4^+^ T cells; CD8: CD8^+^ T cells; DC: dendritic cells; M1: classically activated macrophages; M2: alternatively activated macrophages; Mono: monocytes; Neu: neutrophils; NK: natural killer cells; T: T cells; Treg: regulatory T cells

We then developed a deconvolution algorithm to estimate the absolute proportions (i.e. cell fractions referred to the total cells in the sample under investigation) of ten different immune cell types from bulk RNA-seq data. quanTIseq performs deconvolution using constrained least squares regression^18^ to force the cell fractions to be non-negative and their sum not to exceed 1. By allowing this sum to be lower than 1, quanTIseq estimates also the proportion of uncharacterized cells (referred to as “other” cells from here on), namely cells that are present in the cell mixture of interest but not represented in the signature matrix (e.g. cancer cells). After regression, quanTIseq normalizes the immune cell fractions by a scaling factor in order to correct for differences in total mRNA content per cell. The deconvolution of closely related cell types (e.g. T_reg_ cells and non-regulatory CD4^+^ T cells) is inherently hampered by the high correlation of their expression signatures (multi collinearity) and can result in the underestimation or “dropout” of low-abundance T_reg_ cells^15^. As there is currently no consensus on whether regularization methods can overcome multi collinearity in regression-based deconvolution^19,20^, we adopted an heuristic strategy to specifically address the issue of T_reg_ cell dropouts. Further details on quanTIseq algorithm are reported in the **Methods** section.

Deconvolution methods usually take as input a matrix summarizing the gene expression levels of the mixtures of interest^13^ computed from raw expression data. These data can be profoundly different from the signature matrix used for deconvolution, both in terms of gene annotation and normalization of gene expression values. To avoid issues arising from missing signature genes and different data-normalization procedures, quanTIseq implements a full pipeline for the analysis of raw RNA-seq data that builds the mixture matrix using the same approach employed for the signature matrix (described in **Methods**). The quanTIseq pipeline consists of three analytical steps, as depicted in Figure 1c: 1) Pre-processing of raw RNA-seq reads (single- or paired-ends) to remove adapter sequences, trim low-quality read ends, crop long reads to a maximum length, and remove short reads; 2) Quantification of gene expression as transcripts per millions (TPM)^21^ – which are suitable for expression deconvolution based on linear regression^22^ – and raw counts; 3) Expression normalization, gene re-annotation, and deconvolution of cell fractions. A unique feature of quanTIseq is the possibility to perform *in silico* multiplexed immunoprofiling by complementing the deconvolution results with information from image analysis of IHC, IF, or H&E tissue slides. If total cell densities estimated from images are available, they are used by quanTIseq to scale the fractions of all the deconvoluted immune cell types to cell densities (step 3 in Figure 1c).

quanTIseq analytical pipeline is embedded in a Docker image (https://www.docker.com) that simplifies the installation and usage of all tools and dependencies, thereby standardizing data analysis and making it easily accessible by a broader audience. quanTIseq is available at: http://icbi.i-med.ac.at/software/quantiseq/doc/index.html.

### Validation of quanTIseq using simulated RNA-seq data and published data sets

To benchmark our algorithm on well-defined cell mixtures, we simulated 1,700 RNA-seq data sets of human breast tumors characterized by different immune infiltrate scenarios. The data were generated by mixing different proportions of RNA-seq reads from tumor and immune cells and by simulating different sequencing depths (details in **Methods**). In order not to use the same data for the mixture and signature matrix in the benchmarking, we adopted a leave-K-out cross-validation approach. Briefly, for each simulated mixture to be deconvoluted, a signature matrix was built excluding the *K* RNA-seq data sets included in the simulated mixture. quanTIseq obtained a high correlation between the true and the estimated fractions and accurately quantified tumor content, measured by the fraction of “other” cells (**Supplementary Figure 1**).

We then validated quanTIseq using experimental data from a previous study^23^, in which peripheral blood mononuclear cell (PBMC) mixtures were subjected to both RNA-seq and flow cytometry. A high accuracy of the quanTIseq estimates was also observed for this data set (Figure 1d and **Supplementary Figure 2**). Additionally, we tested quanTIseq on two published microarray data sets used to validate previous deconvolution methods^15,24^. Although quanTIseq is designed for RNA-seq data and might show lower accuracy on pre-computed expression data due to the lack of important signature genes and due to the different dynamic range of hybridization-based and RNA-seq technologies, there was high concordance also with these data sets (**Supplementary Figures 3** and **4**).

We then validated quanTIseq using over 8,000 TCGA samples across 19 solid malignancies. As no gold-standard measures were available for these samples, we considered previous estimates of lymphocytic infiltration^25^ and tumor purity^26^ available for a subset of the TCGA patients to further assess the validity of quanTIseq results. First, we compared the fraction of lymphocytes estimated by quanTIseq, computed by summing up the cell fractions of B cells, NK cells, CD4^+^ and CD8^+^ T cells, and T_reg_ cells, with the “lymphocyte score”, a semi-quantitative measure of the number of tumor-infiltrating lymphocytes estimated previously from H&E-stained section slides of melanoma tumors (n=468)^25^. Although the two approaches were based on different features of the immune contexture, e. molecular vs. morphological, and sequencing data and images are usually generated from different tumor portions, their estimates showed a high agreement (**Supplementary Figure 5a**). Second, we considered TCGA tumor purity values estimated in a previous work with a consensus approach integrating four computational methods based on RNA-seq, methylation, and mutational data^26^. We compared these purity values with the fraction of “other” cells inferred by quanTIseq for all cancer types for which both estimates were available for at least 100 patients. Although the fraction of “other” cells does not directly represent tumor purity as it can include different cell types (e.g. stromal cells), we reasoned that a large proportion of these cells are tumor cells and therefore a positive correlation between these two variables in solid tumors should be expected. Indeed, the fraction of “other” cells estimated by quanTIseq had a significant positive correlation with tumor purity in all cancer types, with a correlation ranging from 0.29 in glioblastoma (GBM) to 0.72 of skin cutaneous melanoma (SKCM) (**Supplementary Figure 5b**).

### Validation of quanTIseq with flow cytometry immunoprofiling and IHC/IF data

Comparison of different deconvolution methods is difficult since the performance strongly depends on the data type and the data pre-processing, as well as on the number and type of immune cells considered (e.g. rare and similar cell types, considered by some methods but not by others, are more difficult to quantify). We therefore initiated extensive validation study and generated data sets which can be also a valuable resource for future, independent benchmarking studies.

As most of the validation data sets available in the literature are based on microarray data or consider a limited number of phenotypes, we generated RNA-seq and flow cytometry data from mixtures of peripheral blood immune cells collected from nine healthy donors. The mixtures were generated by admixing low fractions of polymorphonuclear (PMN) cells with PBMC extracted from the same donor samples (see **Methods**). Flow cytometry was used to quantify all the immune sub-populations considered by quanTIseq except macrophages, which are not present in blood. Comparison of the quanTIseq estimates with the flow cytometry cell fractions showed a high correlation for all the single cell types (Figure 1e and **Supplementary Figure 6**) and an overall correlation of 0.87. In particular, quanTIseq accurately quantified closely related cell types like non-regulatory CD4^+^ T and T_reg_ cells, as well as low-abundance dendritic cells (**Supplementary Figure 6**).

Finally, we validated quanTIseq using three independent cancer data sets. The first data set was generated from 70 tumor samples collected from melanoma patients. We carried out RNA-seq and, wherever possible, IHC staining for CD8^+^, CD4^+^ or FOXP3^+^ cells from consecutive whole-tissue slides. To quantify specific immune cells from the scanned images, we developed an analysis pipeline (available at https://github.com/mui-icbi/IHCount) to perform semi-automatic cell counting. The second data set was generated in an analogous manner using eight lung cancer samples and IHC images stained for CD8^+^ and CD4^+^ T cells. Example images for these two data sets are shown in Figures 2a and **c**. The third data set was generated from tumor samples of ten CRC patients. RNA-seq data, IF-stained slides for CD8^+^ T cells, and IHC slides for CD4^+^ T and T_reg_ cells were generated. Cell densities were then quantified with proprietary software for automated quantitative pathology (details in **Methods**). For one CRC sample, tissue integrity was compromised during the staining procedure and no counts for CD4^+^ T cells could be obtained. The cell fractions obtained with quanTIseq correlated with the respective image cell densities for the melanoma (Figure 2b), lung cancer (Figure 2d), and CRC cohort (Figure 2e). Pearson’s correlation ranged in 0.32-0.75, 0.55-0.91, and 0.58-0.90, respectively. The absolute proportions of the cytotoxic CD8^+^ T estimated by quanTIseq ranged between 0% and ~10%, but did not exceed 3% in the lung cohort. The lower agreement between the deconvolution estimates and cell densities for T_reg_ cells in the melanoma cohort is likely due to the low cell abundances and limited sample size. Overall, the good concordance between the deconvolution results and the gold standard IHC/IF measurements endorses the validity of quanTIseq estimates of tumor-infiltrating immune cells.

**Figure 2:**
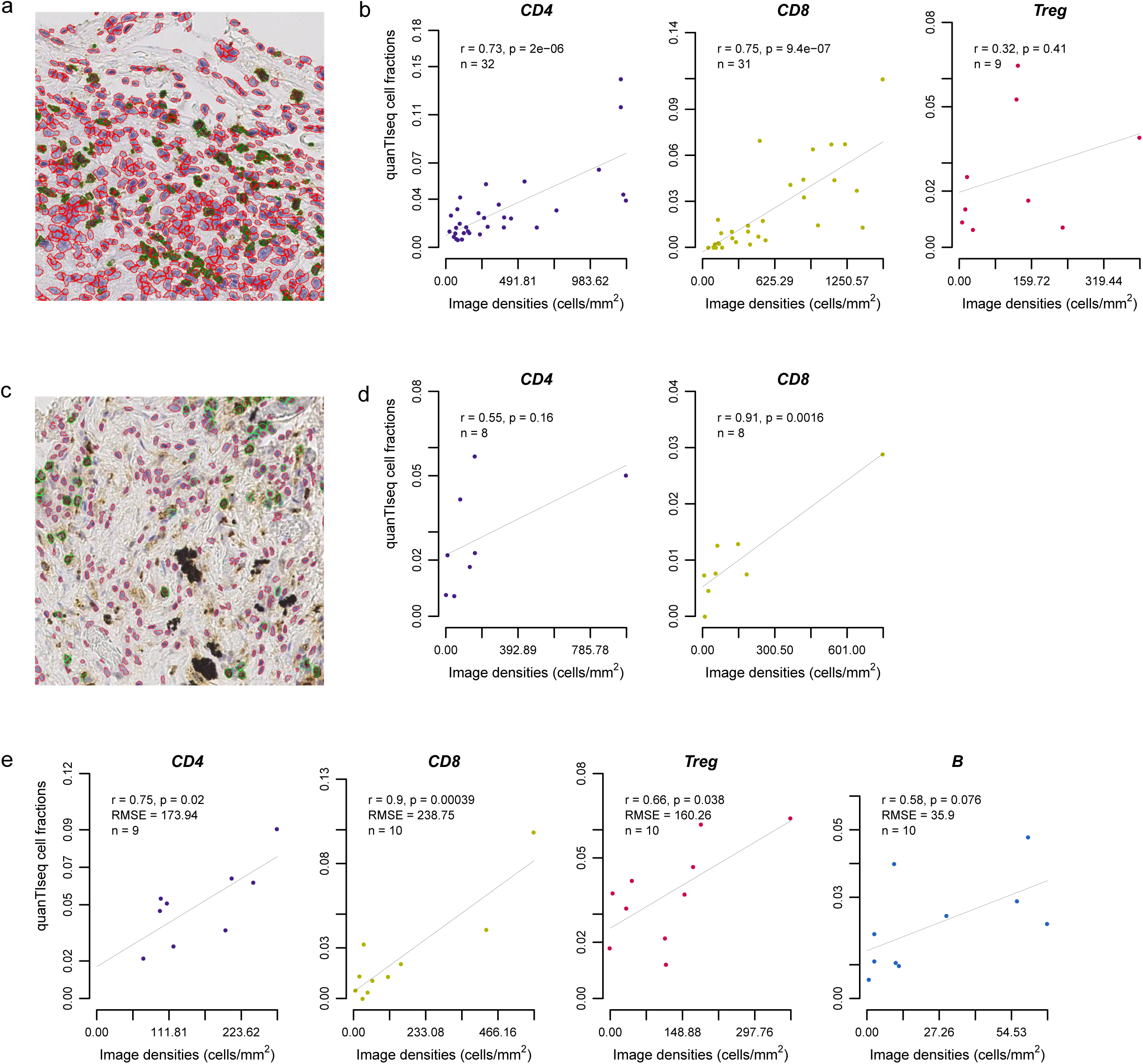
Validation of quanTIseq using tumor RNA-seq data and IF/IHC images. Representative immunohistochemistry (IHC) images from the melanoma **(a)** and lung cancer **(c)** cohorts analyzed with IHCount (magnification 20x, resolution 20000 pixels/cm). The identified nucleated (red) and CD8^+^ cells (green) are highlighted. Comparison of the cell fractions inferred for the melanoma **(b)**, lung cancer **(d)**, and colorectal cancer **(e)** patients. Deconvolution performance was assessed with Pearson’s correlation (r) considering the cells per mm^2^ computed from the IHC or immunofluorescence (IF) images as ground truth. The line represents the linear fit. B: B cells. CD4: total CD4^+^ T cells (including also CD4^+^ regulatory T cells); CD8: CD8^+^ T cells; T: Treg: regulatory T cells.

### Activation of the CXCL9/CXCR3 axis is associated with immune infiltrates in solid cancers

A comprehensive inventory of molecular determinants that shape the immune contexture have not been fully elucidated. In an attempt to identify promising candidates, we examined the association between the immune contexture and a set of features describing the genotypes of human cancers. For this purpose, we used quanTIseq to reconstruct the immune contexture of solid tumors from TCGA RNA-seq data of more than 8,000 TCGA samples across 19 solid malignancies and we assessed the correlation between absolute cell proportions and different genomic features: mutational load, neoantigen load, tumor heterogeneity, and fractions of mutations with clonal and subclonal origin. Surprisingly, there was either low or no correlation between these genetic correlates and the abundances of tumor-infiltrating immune cells (**Supplementary Figure 7**). Moreover, the overall lymphocytic infiltration and the sum of all adaptive or innate immune cell fractions were only weakly associated with the mutational features in our pan-cancer and cancer-specific assessments.

We have previously used biomolecular-network reconstruction to identify T-cell homing factors associated with survival in CRC and pinpointed specific chemokines (CX3CL1, CXCL9, CXCL10) and adhesion molecules (ICAM1, VCAM1, MADCAM1) associated with high densities of intratumoral T cell subsets^27^. Therefore, we assessed the association between the expression of relevant chemokines, chemokine receptors, and adhesion molecules and the abundances of individual immune cell types (**Supplementary Figure 8**). We observed a high correlation between CD8^+^ T cell fractions and the expression of CXCL9 (Figure 3a) and CXCL10 (**Supplementary Figure 8a**) chemokines, as well as with the chemokine receptor CXCR3 (**Supplementary Figure 8b**). The CXCL9/CXCR3 axis regulates immune cell migration, differentiation, and activation and is therefore an important target for cancer therapy^28^.

**Figure 3:**
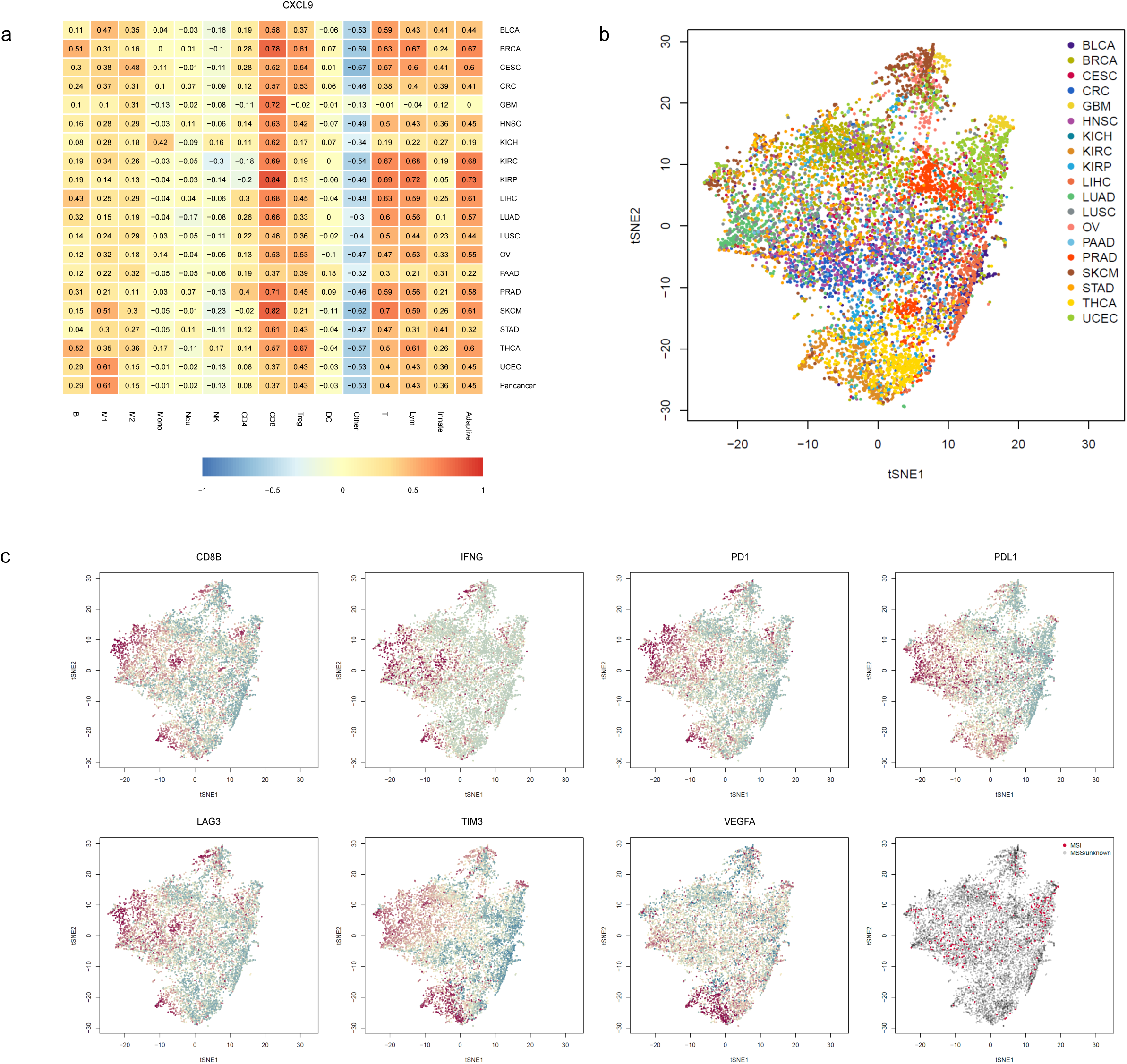
quanTIseq analysis of RNA-seq data from 19 TCGA solid cancers. **(a)** Pearson’s correlation between the cell proportions estimated by quanTIseq and the expression in TPM of the CXCL9 chemokine. t-SNE plot of the immune contextures of 8,243 TCGA cancer patients, colored by: **(b)** cancer type or **(c)** expression of immune-related genes and microsatellite instability state (MSI: microsatellite instable; MSS: microsatellite stable). Adaptive: total adaptive immune cells; B: B cells; CD4: total CD4^+^ T cells (including also CD4^+^ regulatory T cells); CD8: CD8^+^ T cells; DC: dendritic cells; Innate: total innate immune cells; Lym: total lymphocytes; M1: classically activated macrophages; M2: alternatively activated macrophages; Mono: monocytes; MSI: microsatellite instable; MSS: microsatellite stable; Neu: neutrophils; NK: natural killer cells; Other: uncharacterized cells; T: T cells; Treg: regulatory T cells.

In summary, our results obtained using quanTISeq on bulk RNA-seq data from the TCGA support the notion that the activation of the CXCR3/CXCL9 axis, rather than the genotype of the tumor, is associated with intratumoral cytotoxic T cells infiltration, and challenges previous notion that the mutational burden is associated with an increased infiltration of immune cells^29^.

### Pan-cancer analysis reveals highly heterogeneous immune contextures within and across solid cancers

We have previously shown that mutation and neoantigen profiles are highly heterogeneous on a sample by sample basis, being mostly characterized by passenger alterations that are rarely shared between patients^11^. However, despite this huge variability in their genotypes, tumors present common transcriptional signatures describing few molecular subtypes. For instance, analyses of a large number of samples identified four CRC subtypes with clear biological interpretability, called consensus molecular subtypes (CMS) ^30^. Similarly, the immune profiles of human cancers seem can be grouped into three major phenotypes, which are associated to the response to PD1/PDL1 blockade: immune-inflamed, immune excluded, and immune desert^2^. Hence, we hypothesized that despite the genetic heterogeneity human tumors converge to a limited number of immunological states quantified by the immune contextures. To test this hypothesis we used dimensionality reduction based on the t-Distributed Stochastic Neighbor Embedding (t-SNE)^31^ approach to visualize the 8,243 immune contextures reconstructed by quanTIseq across 19 TCGA solid cancers (Figure 3b). Most of the cancer types did not create clearly distinct clusters, indicating highly heterogeneous immune contextures within and across cancers. Although some weak clusterization was visible for subsets of melanoma (SKCM), thyroid cancer (THCA), uterine cancer (UCEC), breast cancer (BRCA), and lung adenocarcinoma (LUAD) patients, a large overlap is seen for most of the cancer types. Visualization of gene expression (Figure 3c) revealed two major clusters that might identify patients characterized by a high infiltration of cytotoxic CD8^+^ T cells typical of the inflamed phenotype (left cluster with high CD8B expression), opposed to the immune-desert phenotype (right cluster with low CD8B expression)^2^. The inflamed phenotype was further associated with high expression of interferon gamma (IFNG), as well as with up-regulation of immune checkpoints like PD1 and PDL1 and exhaustion markers like LAG3 and TIM3. Intriguingly, the plot also shows a cluster of patients characterized by high CD8B and VEGFA expression (bottom cluster), which might correspond to an immune excluded phenotype^2^.

Based on the results of a recent clinical study^32^, cancers with microsatellite instability (MSI) including CRC, uterine cancer, and ovarian cancer can be now treated with PD1 blockers. We therefore analyzed the immune contextures of MSI cancers from the TCGA cohorts (Figure 3c). Similarly to the pan-cancer analyses, we found no distinct clusters also for this subgroup. Compared to their microsatellite stable (MSS) counterparts, MSI cancers were characterized by a significantly lower infiltration of M2 macrophages (p=5.03·10^-8^) and neutrophils (p=1.28·10^-17^), and by a significantly higher infiltration of M1 macrophages (p=3.66·10^-3^), NK cells (p=5.76·10^-7^), CD8^+^ T cells (p=1.75·10^-4^), T_reg_ cells (p=1.34·10^-3^), and dendritic cells (p=3.67·10^-3^).

In summary, we could show that for human solid tumors neither the classification according to the mutational load (MSI vs MSS) nor the classification according to the anatomical site converges to a limited number of immunological states quantified by the immune contextures. However, it appears that some cancer subtypes exhibit similar immune contextures associated with specific genotypes as recently shown by us ^10,11^ and others ^29^.

### Deconvolution-based immune scores are associated with survival in solid cancers

The immunoscore, a scoring system defined to quantify the immune infiltrates from tumor imaging data, has been demonstrated to be a prognostic marker superior to the TNM staging system in CRC^33^. The immunoscore is based on the enumeration of two lymphocyte populations (CD3^+^ and CD8^+^) in the tumor core and invasive margin, and it can assume values from 0, when low densities of both cell types are found in both regions, to 4, when high densities are found in both regions. Recently, it was shown that the immunoscore as well as a newly proposed T- and B-cell score (TB score) were the strongest predictors of disease-free survival and overall survival in CRC with metastatic lesions^34^. To test whether the deconvolution-based immunoscore and TB score are associated with survival in solid cancers, we defined two modified scores based on the absolute fractions of the respective cell types deconvoluted by quanTIseq (see **Methods**).

The results of the survival analysis using the computed TCGA cell fractions showed the prognostic value of the deconvolution-based immunoscore and TB cell score in five and six solid cancers, respectively, namely BRCA, cervical squamous cell carcinoma (CESC), head and neck cancer (HNSC), SKCM, and UCEC, and BRCA, CESC, HNSC, LUAD, and prostate adenocarcinoma (PRAD) (Figure 4). The association was not significant for CRC likely due to the fact that spatial information of the immune cell distribution with respect to the tumor core and invasive margin could not be incorporated.

**Figure 4:**
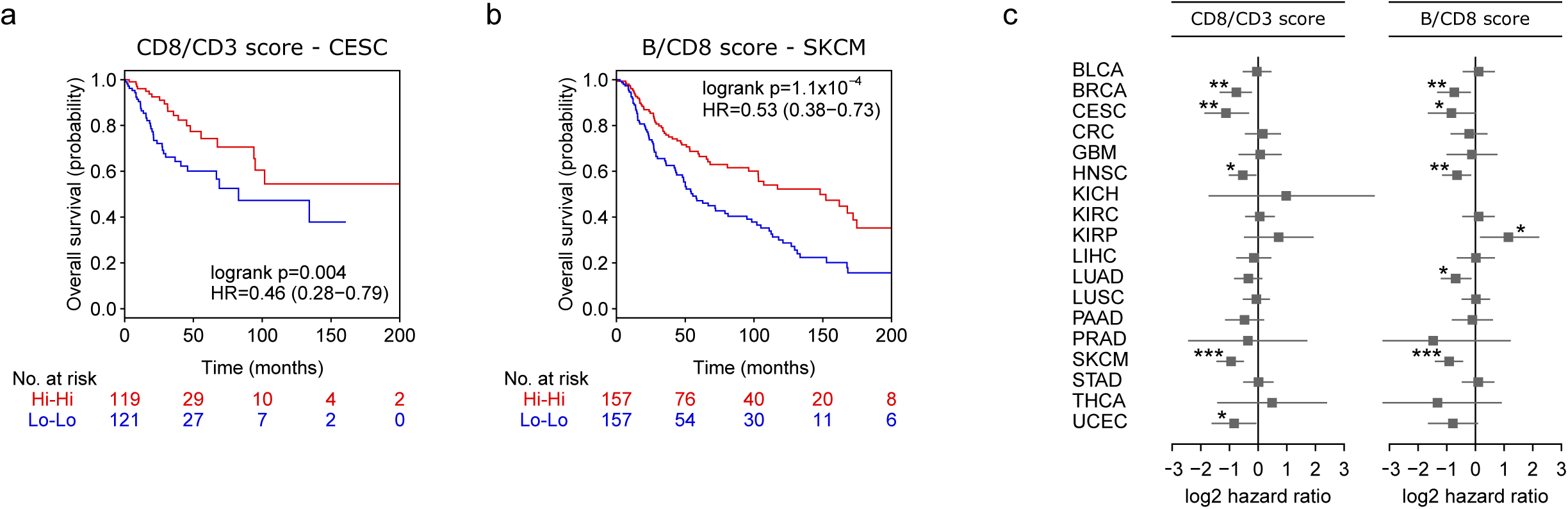
Prognostic value of deconvolution-based Immunoscore and T cell/ B cell score in solid cancers. Kaplan-Meier plots showing the survival of the Hi-Hi and Lo-Lo classes defined considering the deconvolution-based Immunoscore computed for cervical endometrial cancer (CESC) patients **(a)** and the TB score computed for melanoma (SKCM) patients **(b)**. The p-value of the log-rank test, hazard ratio (HR) with 5% confidence intervals, and number of patients at risk at the respective time points are reported. **(c)** Results of the overall survival analysis across 19 TCGA solid cancers. Log_2_ hazard ratio and its 95% confidence interval are visualized for the deconvolution-based Immunoscore and TB score as forest plots. Significant p-values are indicated as: * * * p<0.001,* * 0.001≤p<0.01, and * 0.01≤p<0.05.

**Figure 5:**
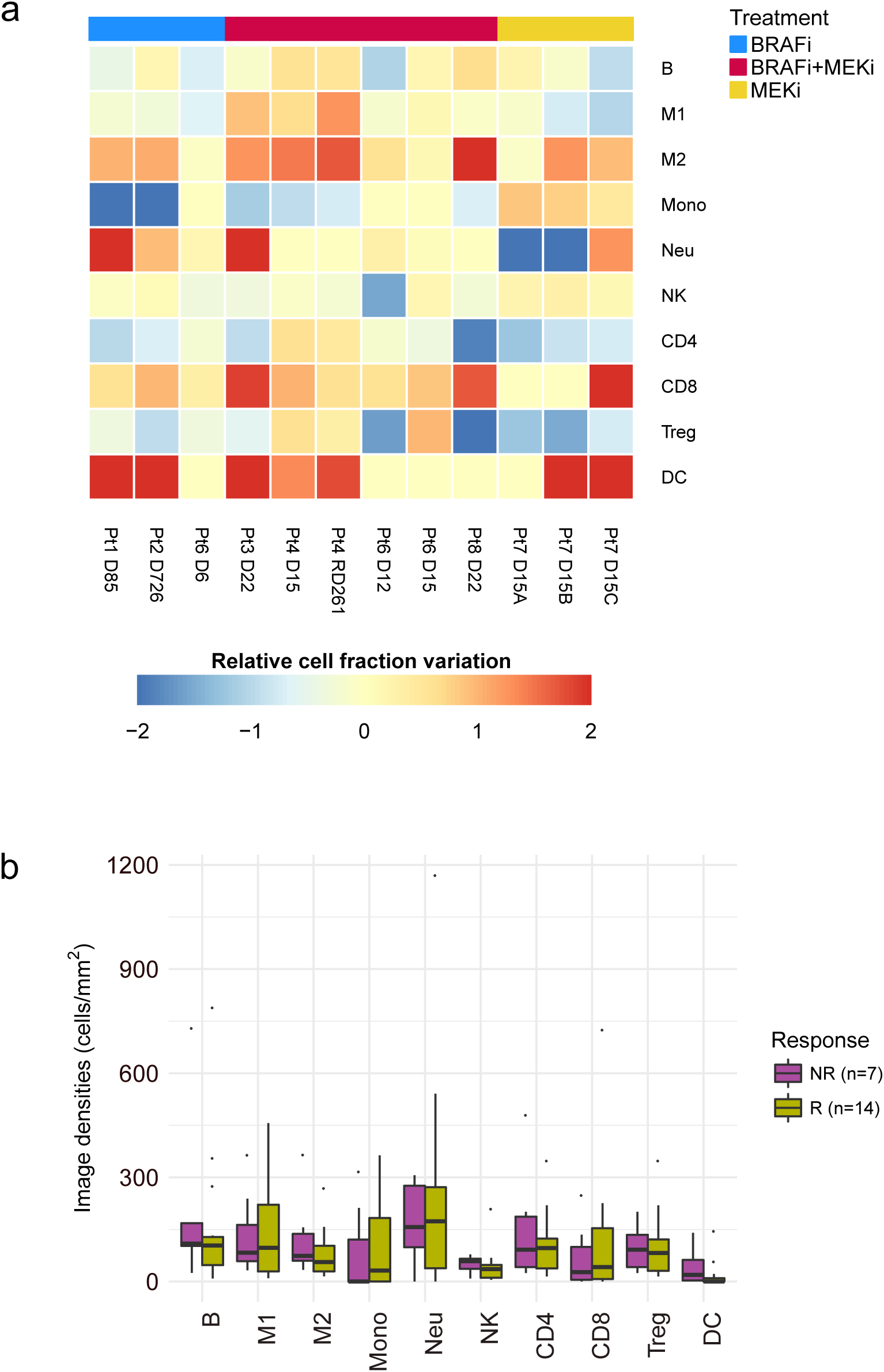
Pharmacological modulation of immune contexture and response to checkpoint blockers. **(a)** Changes in the immune contexture of melanoma tumors after treatment with BRAF and/or MEK inhibitors, measured as the ratio between the difference and the mean of the post- and pre-treatment immune cell fractions. **(b)** Immune cell fractions from melanoma patients stratified as responders (R) and non-responders (NR). B: B cells; CD4: total CD4^+^ T cells (including also CD4^+^ regulatory T cells); CD8: CD8^+^ T cells; DC: dendritic cells; M1: classically activated macrophages; M2: alternatively activated macrophages; Mono: monocytes; Neu: neutrophils; NK: natural killer cells; T: Treg: regulatory T cells; Other: other uncharacterized cells.

All quanTIseq results of the TCGA analysis have been deposited in The Cancer Immunome Atlas (https://tcia.at)^11^ to make them available to the scientific community and facilitate the generation of testable hypotheses.

### Pharmacological modulation of the tumor immune contexture

Beyond the extraction of prognostic markers, there is an urgent need to identify predictive markers for cancer immunotherapy with immune checkpoint blockers, as well as to determine the immunological effects of targeted agents^6^. We therefore used quanTIseq to investigate the pharmacological effects of targeted drugs on the immune contexture. We analyzed recently published RNA-seq data set from pre- and on-treatment tumor biopsies from seven melanoma patients treated with a BRAF inhibitor, MEK inhibitor, or a combination thereof^35^. quanTIseq deconvolution results showed large pharmacological remodeling of the immune contexture (Figure 5a). Changes included a significant increase in dendritic cell fractions after treatment (p=0.043) and, to a lesser extent, an infiltration of CD8^+^ T cells (p=0.19) and M2 macrophages (p=0.07). Thus, BRAF and MEK inhibitors induce profound changes of the immune contexture. However, our analysis showed also patient-specific effects, further highlighting the need to develop immuno-oncology treatment strategies tailored to the individual patient.

Finally, in order to show the value of quanTIseq for informing cancer immunotherapy, we analyzed 21 pre-treatment samples from melanoma patients treated with anti-PD1 antibodies (nivolumab, pembrolizumab) and quantified the immune contexture using bulk RNA-seq data and H&E-stained slides. We first carried out deconvolution using RNA-seq data and then scaled the fractions using cells densities extracted from images to perform *in silico* multiplexed immunodetection. The cell densities of the ten immune cell types showed large heterogeneity across the patients and differences between responders and non-responders. For example, the densities of M1 macrophages as well as of CD4^+^ and CD8^+^ T cells were increased in responders compared to non-responders, although differences were not statistically significant (p>0.09) likely due to the limited number of samples, (Figure 5b). Further work with larger number of samples is necessary to determine which immune cell type fractions or combined scores have predictive power for the response to therapy with immune checkpoint blockers.

## DISCUSSION

We developed quanTIseq, a computational pipeline for the analysis of raw RNA-seq and tissue imaging data that quantifies the absolute fractions and densities of ten different immune cell types relevant for cancer immunology. Unlike previous approaches, quanTIseq is specifically designed for RNA-seq data, which is the current reference technology for high-throughput quantification of gene expression^36^. To simplify data analysis and avoid inconsistencies between the mixture and the signature matrix, we designed quanTIseq as a complete analytical pipeline that performs preprocessing of raw RNA-seq data, gene expression quantification and normalization, gene reannotation, and estimation of cell fractions and densities. The results of our extensive validation using RNA-seq data from simulations, previous studies, blood cell mixtures, and three cancer patient cohorts, demonstrate that quanTIseq can faithfully and quantitatively infer immune-cell fractions from bulk RNA-seq data. Additionally, application of the method to publicly available data as well as data generated in this study revealed several important biological insights.

First, by analyzing more than 8,000 TCGA samples, we showed that genomic features like mutational and neoantigen load, tumor heterogeneity, and proportion of clonal and subclonal mutations are only weakly associated with CD8^+^ T cell fractions. In contrast, the activation of the CXCL9/CXCR3 axis is more strongly associated with CD8^+^ T cell infiltration in solid tumors, supporting the notion that CD8^+^ T cells expressing the chemokine receptor CXCR3 can migrate into tumors following CXCL9 gradients^37^. This finding has an important implication as it suggests that pharmacological modulation of the CXCL9/CXCR3 axis could be a therapeutic strategy to boost T cell recruitment, thereby making also the immune-desert tumors^2^ amenable to cancer immunotherapy. For instance, epigenetic reprogramming of genes expressing T helper (T_H_)-1 chemokines like CXCL9 and CXCL11 might increase CD8^+^ T cell infiltration into the tumor bed^37^.

Second, our results indicate that the immune contexture is highly heterogeneous across and within solid cancers. This could partly explain the fact that the beneficial effects of cancer immunotherapy are observed only in a small fraction of patients. Furthermore, while the classification of common cancers into the three major immunophenotypes, namely immune inflamed, immune excluded and immune desert, is conceptually appealing, it might not be sufficient to stratify the patients and thereby inform cancer immunotherapy. Our data suggest that the immune contexture and, hence, the immunophenotypes, represent rather a continuous then a discrete variable, making it difficult to define cutoffs for precise stratification.

Third, the analysis with the deconvolution-based immunoscore and TB score supports the notion that combinations of different immunological features can have a stronger prognostic power than single markers. The lack of a significant prognostic value for some indications might be due to both, biological and technical reasons. For example analyses of 25.000 samples showed remarkable degree of heterogeneity of the immune infiltrates on the prognosis across distinct organ-specific malignancies ^2^, suggesting that the cellular context is of utmost importance. Moreover, the high heterogeneity of the TCGA cohorts with respect to treatment and staging could be a possible confounding factor. Lastly, as we have previously shown not only the density but also the spatial localization of the infiltrating immune cells plays a major role for the prognosis of tumor recurrence ^3^. Enumeration of the immune cells in the core of the tumor and at the invasive margin markedly enhances the performance of the immunoscore. However, including this type of spatial information from the available TCGA images is challenging due to the limited performance of fully automated image analyses. Spatial lymphocytic patters obtained using recent developments of deep learning tools^29,38^ might provide this missing information.

Forth, quanTIseq analysis of the transcriptomes of patients treated with kinase inhibitors demonstrates profound pharmacological remodeling of the immune contexture. The immunological effects of conventional and targeted therapies came only recently into focus, fostering numerous clinical trials on combinatorial regimens of checkpoint blockers and targeted agents^39^. As bulk RNA-seq is now widely applied to profile fresh-frozen and archived tumor specimens, quanTIseq can be applied to effectively mine these data. Specifically, quanTIseq can be used to quantify the tumor immune contexture from large collections of formalin-fixed paraffin-embedded (FFPE) samples in order to identify immunogenic effects of conventional and targeted drugs and hereby gain mechanistic rationale for the design of combination therapies.

Finally, our analysis of the baseline transcriptomics profiles of patients treated with anti-PD1 antibodies, although limited by the sample size, shows the potential of quanTIseq for the extraction of immunological features that, alone or in combination, might predict the response to checkpoint blockade. As more and more RNA-seq data sets from pre- and post-treatment samples of patients treated with checkpoint blockers will become available, we envision that quanTIseq will represent a useful resource not only to extract candidate predictive markers, but also to monitor the reshaping effect of immunotherapy on the tumor immune contexture.

We envision three lines of improvements of quanTIseq. First, as the transcriptomes of other non-malignant cell types from the tumor microenvironment will become available using bulk RNA-seq or single-cell RNA-seq, quanTIseq signature matrix can be extended to other cell types (e.g. cancer-associated fibroblasts) and optimized for specific cancer types. Second, spatial information on the localization of the infiltrating immune cells, i.e. localization in the center of the tumor and at the invasive margin can be incorporated using annotation by a pathologist from images of H&E-stained slides. Finally, complementary information on the functional orientation of the infiltrating immune cells, including T cell anergy, exhaustion, or differentiation stage, can be derived from bulk RNA-seq data and included into the algorithm. However, since these functional states are not precisely defined in terms of unique expression signatures, a community-based consensus is required in order to include this type of information.

In summary, we developed and thoroughly validated quanTIseq, a method for the quantification of the immune contexture using bulk RNA-seq data and histological images. Application of the tool to analyze thousands of archived samples from patients treated with conventional and targeted drugs revealed molecular and pharmacological modulators of the tumor immune contexture. Hence, by analyzing carefully selected and well annotated archived samples, our method holds promise to derive mechanistic rationale for the design of combination therapies. While quanTIseq represents an important contribution to the computational toolbox for dissecting tumor-immune cell interactions from RNA-seq data^13^, we envision that it can be also applied to study autoimmune, inflammatory, and infectious diseases.

## METHODS

### Collection of RNA-seq data from immune cell types and tumor cell lines

To build the signature matrix, we collected 51 data sets generated from paired-end Illumina RNA-seq of blood-derived immune cells (**Supplementary Table 1**). In addition, we downloaded from the Cancer Genomics Hub (CGHub, https://cghub.ucsc.edu, accessed on February 2016) RNA-seq data from a breast (G41726.MCF7.5) and a colorectal (G27202.SW480.1) cancer cell line. BAM files of mapped reads gathered from to the CGHub were converted to FASTQ with samtools^40^, whereas SRA files downloaded from the Sequence Read Archive (SRA, https://www.ncbi.nlm.nih.gov/sra/) were converted to FASTQ with the “fastq-dump” function of the SRA Toolkit.

### RNA-seq data pre-processing

FASTQ files of RNA-seq reads were pre-processed with Trimmomatic^41^ to remove adapter sequences and read ends with Phred quality scores lower than 20, to discard reads shorter than 36 bp, and to trim long reads to a maximum length of 50 bp. This analysis is implemented in the “Preprocessing” module of quanTIseq (step 1 in Figure 1c), which also allows selecting different parameters for data preprocessing.

### Quantification of gene expression and normalization

The pre-processed RNA-seq reads were analyzed with Kallisto^42^ to generate gene counts and transcripts per millions (TPM) using the “hg19_M_rCRS” human reference. For single-end data, the following Kallisto options were used: “--single -l 50 -s 20”. After gene expression quantification, gene names were re-annotated to updated gene symbols defined by the HUGO Gene Nomenclature Committee (http://www.genenames.org, annotations downloaded on April, 2017). In case of duplicates, the median expression per gene symbol was considered. The final expression value *x* _*gl*_ for each gene *g* in library *l* was computed from TPM with the following formula:

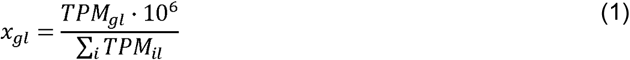

For microarrays data, before the normalization of Equation 1, expression data were transformed from log scale to natural scale (when needed) and quantile-normalized. TPM can be computed from RNA-seq reads with the “Gene Expression Quantification” module of quanTIseq (step 2 in Figure 1c). Gene re-annotation and expression normalization are performed by the quanTIseq “Deconvolution” module before deconvolution (step 3 in Figure 1c) and quantile normalization is performed if the “--arrays” option is set to “TRUE”.

### Generation of the simulated data sets

We simulated RNA-seq data from breast tumors with different purity values and immune infiltrates by mixing pre-processed reads from immune cell types and from a tumor cell line (G41726.MCF7.5) of the RNA-seq compendium. We simulated 100 different immune-cell mixtures by sampling the cell fractions from a uniform distribution in the [0-1] interval. The cell fractions were combined with 11 different tumor purity scenarios: 0:10:100% tumor purity, defined as the fraction of read pairs from the tumor cell line over total read pairs. Each simulated data set consisted of 1 million paired-end reads. In addition, for the data set with 60% purity (which is the minimum value considered by the TCGA consortium for tumor specimen inclusion^26^) we simulated different sequencing depths, namely: 1, 2, 5, 10, 20, 50, and 100 million read pairs. In total, we generated 1,700 simulated RNA-seq data sets. The simulated data are available at: http://icbi.i-med.ac.at/software/quantiseq/doc/index.html.

### Generation of the TIL10 signature matrix

An expression matrix was generated from the compendium of RNA-seq data as described in “RNA-seq data pre-processing” and “Quantification of gene expression and normalization”, and consisted in 19,423 genes and 53 immune- and tumor-cell libraries. From this matrix, we filtered out the genes that were not detected in at least two immune libraries and selected the genes specific for each cell type considering the criteria described in the following. Gene expression is here considered in terms of normalized values *x*_*gl*_ (Equation 1) on a natural scale, if not differently stated.

#### Cell-specific expression

We quantized the expression of each gene into three bins representing low, medium, and high expression, computed as in ^43^. For each immune cell type, we selected the genes having: (i) high quantized expression in all libraries belonging to the considered immune-cell type; and low or medium quantized expression in all other libraries.

#### Expression in tumors

We filtered the signature genes that were highly expressed also in tumor cells by discarding the genes having a median log_2_ expression larger than 7 in all non-hematopoietic cancer cell lines assayed in the Cancer Cell Line Encyclopedia (CCLE)^44^, as done in ^15^. Moreover, RNA-seq data from 8,243 TCGA solid tumors were used to remove genes that provide little support for bulk-tissue deconvolution because their expression in tumor samples is generally low or null. More precisely, we discarded the genes having an average expression across all TCGA samples lower than 1 TPM.

#### Specificity of marker genes

Since signature genes specific for a certain cell type should not be associated to another cell type, we considered a compendium of 489 gene sets specific for 64 cell types recently proposed in ^9^ and removed the signature genes that were listed in a gene set specific for another cell type. CD4^+^ T cell gene sets were not used to filter T_reg_-cell signature genes, as the CD4^+^ T cell population may contain *bona fide* T_reg_-cell expression markers such like the forkhead box P3 (FOXP3).

#### Range of expression

As genes with high expression can bias deconvolution results, we excluded the genes whose expression exceeded the 700 TPM.

#### Correlation with true cell fractions

The 1,700 simulated RNA-seq data sets (see “Generation of the simulated data sets”) were then used to identify the signature genes that provide valuable information over cell fractions and are more robust to the sequencing depth and unknown tumor content. For each cell type, we selected the genes whose expression levels had a correlation with the true cell fractions equal or greater than 0.6.

#### Restricted expression

We considered four external expression data sets from enriched/purified immune cells: two microarray data sets (GEO accession: GSE28490 and GSE2849)^45^, an RNA-seq data set^46^, and a microarray compendium that was used to build the CIBERSORT LM22 signature matrix^15^. All data sets were preprocessed and normalized as explained in the previous paragraphs. For each gene *g* specific for a cell type *c* in the signature matrix, we computed the ratio *R*_*gd*_ between the median expression across all libraries in data set *d* belonging to the cell type *c* and the median expression across all libraries in data set *d* not belonging to the cell type *c*. For each cell type, the top 30 ranked signature genes (or less, when not available) with *median*_*d*_ (*R*_*gd*_)≥ 2 were selected for the final signature matrix. When processing the T_reg_ signature genes, the data sets belonging to CD4^+^ T cells were not considered. T_reg_ signature genes were further filtered with a similar approach, but considering the RNA-seq data of circulating CD4^+^ T and T_reg_ cells from ^47^ and selecting only the genes≥with *median*_*d*_(*R*_*gd*_)≥ 1.

The final signature matrix TIL10 (**Supplementary Table 2**) was built considering the 170 genes satisfying all the criteria reported above. The expression profile of each cell type c was computed as the median of the expression values *x*_*gl*_ over all libraries belonging to that cell type:

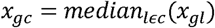

For the analysis of RNA-seq data quanTIseq further reduces this signature matrix by removing a manually-curated list of genes that showed a variable expression in the considered data sets: *CD36, CSTA, NRGN, C5AR2, CEP19, CYP4F3, DOCK5, HAL, LRRK2, LY96, NINJ2, PPP1R3B, TECPR2, TLR1, TLR4, TMEM154, CD248.* This default signature considered by quanTIseq for the analysis of RNA-seq data consist of 153 genes and has a lower condition number then the full TIL10 signature (6.73 compared to 7.45), confirming its higher cell-specificity. We advise using the full TIL10 matrix (-- rmgenes=“none”) for the analysis of microarray data, as they often lack some signature genes, and the reduced matrix (--rmgenes= “default”) for RNA-seq data. Alternatively, the “rmgenes” option allows specifying a custom list of signature genes to be disregarded (see quanTIseq manual).

### Deconvolution

The quanTIseq deconvolution module takes as input:

- A mixture matrix *M*_*gj*_ of expression values over *g* = 1,…,*I* genes and *j* = 1,…, *J* samples;
- A signature matrix *S*_*gc*_ of expression values over *g* = 1,…,*G* signature genes and *c* = 1,…,*C* cell types.

After re-annotation of gene symbols and normalization of the mixture matrix (see “Quantification of gene expression and normalization”), quanTIseq performs deconvolution of the unknown cells fractions *F*_*cj*_ over *C* immune cell types and *J* samples. For each sample *j*, the following system of equations is solved to estimate the cell fractions *F*_*c*_ (the subscript *j* is omitted):

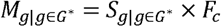

where *G*^*\**^ is the set of signature genes that are present in the mixture matrix. quanTIseq solves this inverse problem using constrained least squares regression, i.e. by minimizing the formula: ‖***S***×*F* - ***M****2*, imposing the constraints:

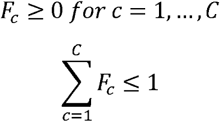

To account for differences in the average mRNA content per cell type, which might otherwise bias deconvolution results^16,48,49^, the estimated cell fractions are normalized by a cell-type-specific scaling factor *n* _*c*_:

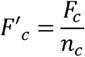

Then, the cell fractions are scaled so to sum up to the original percentage of total cells, as:

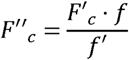

where

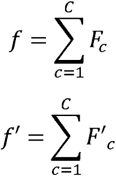

Finally, the proportion of “other” (uncharacterized) cells is estimated as:

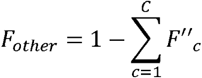

As the population of “other” cells might include different types of malignant and normal cells with various mRNA contents^50^ depending on the sample under investigation, quanTIseq does not scale these estimates. The scaling factors *n*_*c*_ were computed as the median expression of the Proteasome Subunit Beta 2 (PSMB2) housekeeping gene^51^ in the immune cell types of the RNA-seq compendium. In the analysis of the simulated RNA-seq data, where the true fractions represented mRNA fractions and not cell fractions, deconvolution was performed without mRNA-content normalization (**Supplementary Table 4**).

The deconvolution of T_reg_ cells and CD4^+^ T cells is inherently hampered by the high correlation of their expression signatures (namely, *multi-collinearity*^15^) and can result in the underestimation of T_reg_ cells present in low fractions. Thus, we adopted an heuristic strategy to specifically address the issue of T_reg_-cell underestimation. First, quanTIseq estimates the T_reg_ cell fractions 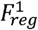 considering all cell types together. Then, for the samples with 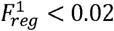 < 0.02, quanTIseq re-estimates the T_reg_ cell fractions 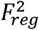 removing from the signature matrix the expression profiles of the CD4^+^ T cells. The final T_reg_ cell fractions are then estimated by averaging the results:

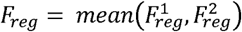

whereas CD4^+^ T cell fractions are scaled to:

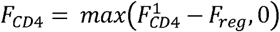

Finally, all cell fractions are normalized to sum up to 1.

The analysis described in this section is implemented in the “Deconvolution” module of quanTIseq (step 3 in Figure 1c).

The full quanTIseq pipeline can be applied to single or multiple samples and can be initiated at any step. For instance, pre-computed expression matrices can be analyzed directly with the deconvolution module (step 3 in Figure 1c), although particular care must be taken when performing data preprocessing and annotation of signature genes.

### Deconvolution of bulk tumor expression data

Aberrant de-methylation and sequence duplication can lead to over-expression of immune signature genes. Tumor RNA-seq data can be analyzed with quanTIseq setting the “--tumor” option to “TRUE”. This setting discards the signature genes whose *log*_*2*_(*x*_*gl*_ + 1) expression in the TCGA RNA-seq data exceed 11 TPM, which are: *NUPR1, CD36, CSTA, HPGD, CFB, ECM1, FCGBP, PLTP, FXYD6, HOPX, SERPING1, ENPP2, GATM, PDPN, ADAM6, FCRLA, SLC1A3.* All tumor data sets presented in this work have been analyzed with this parameter setting (**Supplementary Table 4**).

### Validation data sets

To benchmark quanTIseq, we considered the expression data sets listed in **Supplementary Table 3**, using the options reported in **Supplementary Table 4**. Normalized microarray data were downloaded from the Gene Expression Omnibus (GEO) (https://www.ncbi.nlm.nih.gov/geo) with the GEOquery R package^52^. Probes were mapped to gene symbols with the biomaRt R package^53^. In case of multiple probes mapping to the same gene symbol, the probe with the highest average expression across all samples was selected. Immune cell fractions estimated with flow cytometry, Coulter Counter, or from images of stained tissue slides were used as ground truth to validate quanTIseq. Where necessary, different functional states of an immune cell type were aggregated by summing up the corresponding cell fractions (e.g., for the Newman’s data set ^15^, B cells were quantified summing up the fractions of naïve and memory B cells).

### Generation of flow cytometry and RNA-seq data from blood-derived immune-cell mixtures

Blood samples from healthy human donors were obtained from the Blood Bank Innsbruck under approval of the local ethics committee. Peripheral blood mononuclear cells (PBMC) were isolated from human whole blood by density centrifugation using Lymphocyte Separation Medium (Capricorn, Ebsdorfergrund, Germany). The PBMC fraction was collected and washed three times with Dulbecco’s phosphate buffered saline. To isolate polymorphonuclear (PMN) cells, the cells on top of the erythrocytes were collected and contaminating red blood cells were removed by two rounds of lysis with 0.2% NaCl solution at 4°C. PMN were added to the PBMC fractions in low abundance (3-6% of total cells) and aliquots were taken for RNA extraction and flow cytometry analysis. Total RNA was extracted with the Qiagen RNeasy mini kit (Qiagen GmbH, Hilden, Austria), including on-column DNAse I treatment. INVIEW polyA RNA library preparation, and Illumina 50bp SR sequencing at >60 Million reads per library was obtained from an external provider (GATC biotech, Konstanz, Germany).

The fractions of the following cell types in the immune-cell mixtures were determined by flow cytometry using specific marker combinations: CD4^+^ T cells (CD3^+^CD4^+^), CD8^+^ T cells (CD3^+^CD8^+^), T_reg_ cells (CD3^+^CD4^+^CD25^+^CD127^-^), B cells (CD19^+^), NK cells (CD3^-^CD16^+^CD56^+^), myeloid dendritic cells (Lin^-^ HLA-DR^+^CD11c^+^), monocytes (CD14^+^) and neutrophils (CD15^+^CD16^+^). Labeled antibodies specific for the following antigens were purchased from BD Biosciences (San Jose, CA, USA) and Biolegend (San Diego, CA, USA): CD3 (UCHT1), CD4 (RPA-T4), CD8 (HIT8a), CD11c (3.9), CD14 (M5E2), CD15 (W6D3), CD16 (3G8), CD19 (HIB19), CD20 (2H7), CD25 (BC96), CD56 (B159), CD127 (A019D5), HLA-DR (L243), Lin: CD3, CD14, CD19, CD20, CD56. The measurements were performed on a BD LSRFortessa flow cytometer and the data were evaluated with FlowLogic 7.1 software (Inivai Technologies, Melbourne, Australia).

### Leiden validation data set

Fresh frozen and formalin-fixed material was available from ten colorectal cancer patients. Their usage was approved by the local ethics committee. All the specimens were anonymized and handled according to the ethical guidelines described in the Code for Proper Secondary Use of Human Tissue in the Netherlands of the Dutch Federation of Medical Scientific Societies. RNA was isolated with the NucleoSpin RNA kit (Macherey-Nagel, Düren, Germany) including on-column DNAse I treatment. Library preparation was preceeded by rRNA depletion with the NEBNext rRNA depletion kit (New England Biolabs, MA, USA). PE 150bp sequencing was performed at GenomeScan (Leiden, The Netherlands) on a HiSeq 4000 (Illumina, San Diego CA, USA).

4μm sections of formalin-fixed paraffin-embedded tissues were deparaffinized and underwent heat-mediated antigen retrieval in 10 mmol/L citrate buffer solution (pH 6). Unspecific antibody binding was prevented with the SuperBlock PBS buffer (Thermo Fisher Scientific, Waltham, MA, USA) according to the manufacturer’s instructions. Immunofluorescence detection was performed with the following antibodies: pan-cytokeratin (AE1/AE3, Thermofisher scientific and C11, Cell Signalling Technology), anti-CD3 (D7A6E), and anti-CD8 (4B11, DAKO). Immunofluorescent detection was performed directly and indirectly with Alexa488, Alexa594 and CF555 with an in-house methodology (manuscript in preparation).

For immunohistochemical detection 4μm sections were deparaffinized after which endogenous peroxidase was blocked with a 0.3% hydrogen peroxide/Methanol solution. Following heat mediated antigen retrieval in 10 mmol/L citrate buffer solution (pH 6), overnight labeling was performed with anti-CD4 (EPR68551, Abcam), anti-FOXP3 (236A/E7) and CD20 (L26, Dako) respectively. After washing in PBS, Tissue sections were incubated for one hour with Poly-horseradish peroxidase solution (Immunologic Duiven, The Netherlands) at room temperature. The slides were developed with the DAB+ chromogen (DAKO, Agilent technologies, Santa Clara, Ca, USA) solution and counterstained with Haematoxylin (Thermo Fisher Scientific).

Image analysis for both immunofluorescence and immunohistochemistry was performed with the Vectra 3.0 Automated Quantitative Pathology Imaging System and the inFORM Cell Analysis software (Perkin Elmer, Waltham, MA, USA) including spectral separation of dyes, tissue and cell segmentation, and automated cell counting of immune phenotypes.

### Vanderbilt validation data sets

70 melanoma and 8 lung cancer patient samples were procured based on availability of tissue and were not collected according to a pre-specified power analysis (**Supplementary Table 5**). Included in these, 42 melanoma samples and 7 lung cancer samples were baseline pre-anti-PD1 therapy. Remaining patients were treated with either anti-CTLA-4 alone or combinations of anti-PD-1 and anti-CTLA-4. Finally, 10 samples were obtained from progressing tumors in patients experiencing an initial response. Clinical characteristics and objective response data were obtained by retrospective review of the electronic medical record. Patients were classified in responders (Complete Response and Partial Response) and non-responders (Progressive Disease, Mixed Response, and Stable Disease) according to investigator assessed, RECIST defined responses. All patients provided informed written consent on IRB approved protocols (Vanderbilt IRB # 030220 and 100178).

Total RNA quality was assessed using the 2200 Tapestation (Agilent). At least 20ng of DNase-treated total RNA having at least 30% of the RNA fragments with a size >200 nt (DV200) was used to generate RNA Access libraries (Illumina) following manufacturer’s recommendations. Library quality was assessed using the 2100 Bioanalyzer (Agilent) and libraries were quantitated using KAPA Library Quantification Kits (KAPA Biosystems). Pooled libraries were subjected to 75 bp paired-end sequencing according to the manufacturer’s protocol (Illumina HiSeq3000). Bcl2fastq2 Conversion Software (Illumina) was used to generate de-multiplexed Fastq files.

For FOXP3, CD4 and CD8 IHC staining, slides were placed on a Leica Bond Max IHC stainer. All steps besides dehydration, clearing and coverslipping were performed on the Bond Max. Heat induced antigen retrieval was performed on the Bond Max using their Epitope Retrieval 2 solution for 20 minutes. Slides were incubated with anti-CD4 (PA0427, Leica, Buffalo Grove, IL), FOXP3 (14-4777-82, eBiosciences), or anti-CD8 (MS-457-R7, ThermoScientific, Kalamazoo, MI) for one hour.

### Analysis of Vanderbilt images with IHCount

We considered 75 brightfield immunohistochemistry images from 33 melanoma patients and 16 images from 8 lung cancer patients. However, 3 melanoma patients had to be excluded from the analysis due to the low quality of the staining or lack of tissue. In total, we analyzed 72 CD4, CD8 and FoxP3 stained images from 32 melanoma patients and 16 CD4 and CD8 stained images from 8 lung cancer patients. To quantify both the number of total cells and tumor-infiltrating immune cells from the melanoma and lung cancer images, we implemented a computational workflow, called IHCount, using free open-source software tools. In this workflow different analytical tasks were performed, including: image pre-processing, training of pixel classifiers, image segmentation and analysis, together with cell counting and additional measurements of the tumor-covered area. The methodology of the analysis is described as follows.

For the purpose of initial pre-processing of the IHC images, we used the script collection (bftools) from the consortium of Open Microscopy Environment (OME)^54^. First, the bright-field images were extracted as TIF files with the highest resolution from the image containers, available in Leica (SCN) format. Each of these high-resolution images (0.5 μm/pixel, 20x magnification) was then subdivided into equally sized non-overlapping image tiles (2000×2000 pixels) in order to limit the computational costs of the subsequent analytical tasks. The open-source software Ilastik^55^ and it’s ‘Pixel Classification’ module was used to manually annotate objects of interest in the tile and generate classifiers that distinguish positively stained cells and nuclei from background and stromal tissue. For each sample a set of 3 to 5 representative image tiles was randomly selected for training considering the diverse nature of the obtained images (caused e.g. by presence of artifacts, differences in illumination and staining intensities). As a result we obtained two classifiers, the first one to classify pixels belonging to positively stained cells and the second one for nuclei. Both of them were also trained on background and stromal tissue. The classifiers were subsequently used in a batch process to obtain two sets of so called “probability maps” for each tile of the sample. Both sets were exported as multichannel TIF (32-bit float), where each channel represented the probabilities of one of the given classes (positively stained cells or nuclei, together with stromal tissue and background). The cell segmentation and cell counting as the last steps of the analysis, were performed with the open-source software Cellprofiler^56^. As input files we used the previously generated probability maps together with the original tiles. In order to segment and identify positively stained cells, nuclei and the area of the total tissue, a pipeline of intensity-based operations (IHCount.cppipe) was defined. The overall results for each sample were obtained by summing up the results of the single tiles.

All previously described steps of the analysis were implemented in a python script (runCP.py) and can be run from the command line. The pipeline, together with a description of the workflow, is publicly available at: https://github.com/mui-icbi/IHCount. IHCount results for the Vanderbilt cohorts are reported in **Supplementary Table 5**. Total cell densities per tumor sample to be used to scale quanTIseq cell fractions were estimated as the median number of all cell nuclei per mm^2^ across all images generated from that tumor.

### Computation of the deconvolution-based Immunoscore and TB score from quanTIseq cell fractions

For the calculation of the deconvolution-derived Immunoscore, we considered the fractions of CD8^+^ T cells and CD3^+^ T cells, were the latter was computed as the sum of CD8^+^ T cell, CD4^+^ T cell, and T_reg_ cell fractions. CD3^+^ and CD8^+^ T cell fractions were dichotomized considering their median across all patients, computed separately for each cell type and cancer type, and used to identify two groups of patients: (1) “Lo-Lo” patients, with both CD3^+^ and CD8^+^ T cell fractions lower or equal to the median; (2) “Hi-Hi” patients, with both CD3^+^ and CD8^+^ T cell fractions higher than the median. The “Hi-Hi” and “Lo-Lo” classes for the T- and B-cell (TB score) were derived in an analogous manner, but considering the fractions of B cells and CD8^+^ T cell estimated by quanTIseq.

### t-SNE plots

t-SNE plots of the TCGA solid cancers were generated with “Rtsne” R package. The t-SNE algorithm was run on the immune cell fractions estimated by quanTIseq, excluding the fraction of uncharacterized cells. We retrieved the annotation about miscrosatellite instability from a recent paper^57^, considering both the MSI categories of the TCGA consortium and the MSI/MSS classes predicted at a confidence level of 0.75. Unambiguous predictions were used to identify the MSI or MSS samples, whereas ambiguous predictions (MSI:1 and MSS:1), null predictions (MSI:0 and MSS:0), or unavailable samples were assigned to the “unknown” MSI state. Gene expression represented as z scores of log2(TPM+1). Before plotting, z scores higher than 3 (or lower than −3) were saturated to 3 (or −3).

### Statistical analysis

Correlation between numeric variables was assessed with Pearson’s correlation. The area under the receiver operating characteristic curve (AUROC) for multi-class classification was computed with the “multiclass.roc” function of the pROC R package. Constrained least squares regression was performed with the “lsei” function from the “limSolve” R package. The root-mean-squared error was computed as 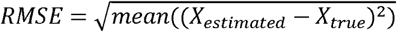. Statistically significant differences between two groups were tested with two-sided Wilcoxon’s test. For comparisons across multiple groups, Kruskal-Wallis test followed by two-sided Dunn’s pairwise post hoc were used. Normality of the data distribution was tested with Shapiro-Wilk test. Overall survival analyses were performed using the R package *survival* on TCGA survival data (‘vital_status’,‘days_to_death’, and ‘days_to_last_followup’). For each cancer type, patients were dichotomized in two groups according to the deconvolution-based Immunoscore or TB score. The Kaplan-Meier estimator was used to generate survival curves and logrank tests (corresponding to two sided z-test) were applied.

## Data availability

The RNA-seq data from blood-derived immune cell mixtures have been deposited in the Gene Expression Omnibus (GEO) under accession number GSE107572. The simulated RNA-seq data sets are available at: http://icbi.i-med.ac.at/software/quantiseq/doc/index.html. All quanTIseq results from TCGA and from the melanoma and lung cancer cohorts have been deposited in The Cancer Immunome Atlas (https://tcia.at)^11^.

## ACKNOWLEDGEMENTS

We would like to thank Dr. Paul Hoertnagl (Innsbruck Blood Bank, Austria) for the collection of blood samples from healthy donors and Dr. Kristen L. Hoek (Vanderbilt Institute for Infection, Immunology and Inflammation, US) for providing access to the flow cytometry data for algorithm validation. This work was supported by the European Union’s Horizon 2020 research and innovation program under grant agreement No. 633592 (project “APERIM: Advanced Bioinformatics Tools for Personalised Cancer Immunotherapy”), by the Austrian Cancer Aid/Tyrol (project No. 17003, “quanTIseq: dissecting the immune contexture of human cancers”), by the Vienna Science and Technology Fund (Project LS16-025), and the Austrian Science Fund (Projects I3291 and I3978 and T 974-B30). The Leiden cohort work was supported by the Fight Colorectal Cancer-Michael’s Mission-AACR Fellowship (2015) and Alpe d’HuZes/KWF Bas Mulder Award (UL2015-7664). The Vanderbilt cohort work was supported by the Vanderbilt-Incyte Research Alliance Program Grant (JMB, DBJ, YX), as well as R00CA181491 (JMB), K23CA204726 (DBJ), and the Breast Cancer Specialized Program of Research Excellence (SPORE) P50 CA098131.

## Author contributions

FF developed quanTIseq and performed deconvolution analyses and method benchmarking. FF and CP implemented quanTIseq software. FF and HH performed the statistical analyses. CM developed IHCount. CM, DR, and ZL analyzed the Vanderbilt images. ZT, FF, AK, GL, WP, DW and SS designed the validation experiment on blood mixtures. GL isolated the blood-derived immune cells and performed the flow cytometry experiment and data analysis. AK performed RNA extraction from the blood-derived immune cell mixtures. DJ contributed to the collection of Vanderbilt patient specimens and clinical annotation. YX and YW oversaw pre-processing and quality control of Vanderbilt RNA-seq data. MES oversaw histology techniques, analysis, and scoring techniques for the Vanderbilt cohorts. MVE and PEG performed IHC data pre-processing for the Vanderbilt cohorts. JB performed Vanderbilt data analysis, integration of techniques, performed and oversaw molecular assays and work involved in sample selection and isolation of nucleic acids. MI and NdM generated RNA-seq and multiplexed IF detection data for the Leiden cohort. FF and ZT wrote the manuscript. All authors read and approved the final manuscript.

